# Spatial heterogeneity in tumor adhesion qualifies collective cell migration

**DOI:** 10.1101/2023.10.05.559967

**Authors:** C V S Prasanna, Mohit Kumar Jolly, Ramray Bhat

## Abstract

Collective cell migration, a canon of most invasive solid tumors, is an emergent property of the interactions between cancer cells and their surrounding extracellular matrix (ECM). However, tumor populations invariably consist of cells expressing variable levels of adhesive proteins that mediate such interactions, disallowing an intuitive understanding of how tumor invasiveness at a multicellular scale is influenced by spatial heterogeneity of cell-cell and cell-ECM adhesion. Here, we have used a Cellular Potts model-based multiscale computational framework that is constructed on the histopathological principles of glandular cancers. In earlier efforts on homogenous cancer cell populations, this framework revealed the relative ranges of interactions, including cell-cell and cell-ECM adhesion that drove collective, dispersed, and mixed multimodal migrations. Here, we constitute a tumor core of two separate cell subsets showing distinct intra-subset cell-cell or cell-ECM adhesion strengths. These two subsets of cells are arranged to varying extents of spatial intermingling, which we call the heterogeneity index (HI). Our simulations show that for a given two intra-subset cell-cell adhesions for two subsets of cells, low and high inter-subset cell adhesion favors migration of high HI and low HI intermingled populations, respectively. In addition, for the most explored values of cell-ECM adhesion strengths, populations with high HI values collectively migrate better than those with lower HI values. We then asked how spatial migration is regulated by progressively intermingled cellular subsets that were epithelial, i.e., showed high cell-cell but poor cell-ECM adhesion, and mesenchymal, i.e., with reversed adhesion strengths to the former. Here too, inter-subset adhesion plays an important role in contextualizing the proportionate relationship between HI and migration. We also observe an exception to this relationship for cases of heterogeneous cell-ECM adhesion where sub-maximal HI patterns with higher outer localization of cells with stronger ECM adhesion collectively migrate better than their relatively higher HI counterparts. Our simulations also reveal how adhesion heterogeneity qualifies migrative dynamics through collective cellular unjamming, when either cell-cell or -ECM adhesion type is varied but incorporates dispersion when both adhesion types are simultaneously altered.

## Introduction

Transformed cells from epithelial tissues and organs migrate from their locus of origin into their surrounding extracellular matrix (ECM) microenvironment in various ways: groups of cells may move together, a process known as collective cell migration (CCM); in contrast, single cells can separate out and migrate independently of each other, which is called dispersed cell migration (DCM) [1]–[3]. Collective cell migration is a complex process that involves coordination by cells across their moving mass, along with cooperative signaling that allows cells to synchronize their actions across the multicellular spatial scale [4], [5]. Although adhesion between moving cancer cells, mediated through cadherin-rich adhesion junctions has been proposed to contribute to collective migration, recent experimental and computational studies suggest this to not be an imperative criterion: non-adherent cancer cell masses may still migrate together, albeit in an uncoordinated manner when allowed to move through a confining ECM space [3], [6]. Interactions with the latter involve both adhesion to ECM proteins and proteolytic or non-proteolytic mechanical effects exerted on them by invading cancer cells [7]. Thus, CCM dynamics is likely influenced by heterogeneous subpopulations of cells with varying adhesive strengths among themselves as well as to the ECM.

Experimental migration studies performed with cancer cell lines or primary cells isolated from patient tumors frequently presume the interactions between cells or with their surrounding ECM to be similar across the collective mass. Although cytological variegation within tumors has long been reported by pathologists, emerging studies on tumor samples establish heterogeneity at multiple levels including at the scale of proteins mediating inter-cellular or cell-ECM adhesion. Alexander and coworkers show for instance that invasive lobular carcinomas thought to be depleted of E-cadherin may instead show cell-variable heterogeneity in its expression [8]. Epigenetic downregulation of E-cadherin driven by the microenvironment may also result in a variegated expression of the protein in ductal breast cancers [9]. Heterogeneity has also been observed experimentally for receptors such as integrins, which mediate adhesion with ECM [10], [11]. Such variation in spatial expression of adhesive proteins suggests concurrent heterogeneity in the force of inter-cellular and cell-ECM adhesion and can be associated with patient survival as well [12].

Heterogeneity in molecular expression within tumors can have marked effects on cellular and multicellular phenotypes. One example is patchy mechanical behavior within cell populations, which is confirmed through their rigorous spatial exploration using atomic force microscopy [13]. The effects of such heterogeneity on cancer cell migration are beginning to be elucidated [14]– [17]. Such patchy behaviors can be quantified via persistence length, as quantified through approaches such as topological data analysis of dynamics of collective migration [18]. Even so, the consequences of variation in the arrangement of adhesive forces across mesoscopic cell scales have rarely been studied. One notable effort by Reher and coworkers uses a cellular automaton approach to show that the dissemination of cells can be mediated through an increase in spatial heterogeneity in cell adhesion and a loss of control on their adhesion receptor concentrations through the local cellular microenvironment [19]. In this paper, we implement a Cellular Potts Model (CPM)-based computational multiscale framework that explores the behavior of a population of cells that are encased within an ECM architecture that is mimetic of the histopathological features of glandular tissues in a precancerous stage. We show how the nature of spatial arrangement in cell-cell or cell-ECM adhesive forces strengths can have significant effects on the dissemination of such tumor populations. Our results are consistent with, and rationalize, the multiple molecular variations associated with epithelial to mesenchymal transition seen in cancers and have implications for the use of biopsies in the diagnosis of invasive cancers.

## Results

### Impact of inter-subset and intra-subset cell-cell adhesion spatial heterogeneity on collective cell migration

We began our study by examining the effects of variation in adhesion strengths on migration in our framework under conditions when such strengths have been kept uniform across the cell population (“homogeneous pattern”) **(Fig 1)**. Our previous work had identified adhesion strengths (we refer to them for the sake of convenience in this paper as “model’’ adhesion strength) at which we had observed CCM [3]. CCM and DCM were assessed as before by measuring the area of the largest connected cluster of cells, and the number of disconnected cells or their clusters, respectively. When cell-cell adhesion strength was halved or doubled, the cell population with homogeneous adhesion pattern showed higher and lower CCM respectively consistent with a frequently observed role in experiments of intercellular homotypic adhesion mediators as suppressors of metastasis [20] (**Fig 1A**; ANOVA p<0.0001). High adhesion to fibrillar ECM such as Collagen I through integrins has been shown to be important for CCM [21], [22]. In support, when the adhesion strength between cancer cells and stromal ECM (in most epithelial tissues, the predominant ECM in stroma is Collagen I) was halved or doubled, the cell population with homogeneous pattern showed lower and higher CCM, respectively **(Fig 1B**; ANOVA p<0.0001). DCM as measured through disconnect in motile cells or clusters in both cases was marginally altered (for one of the subcases only). These results are consistent with experiments that posit an antagonistic interplay between cell-ECM and intercellular adhesive interactions that drive the disaggregation of cellular clusters leading to migration [23].

**Figure 1:**
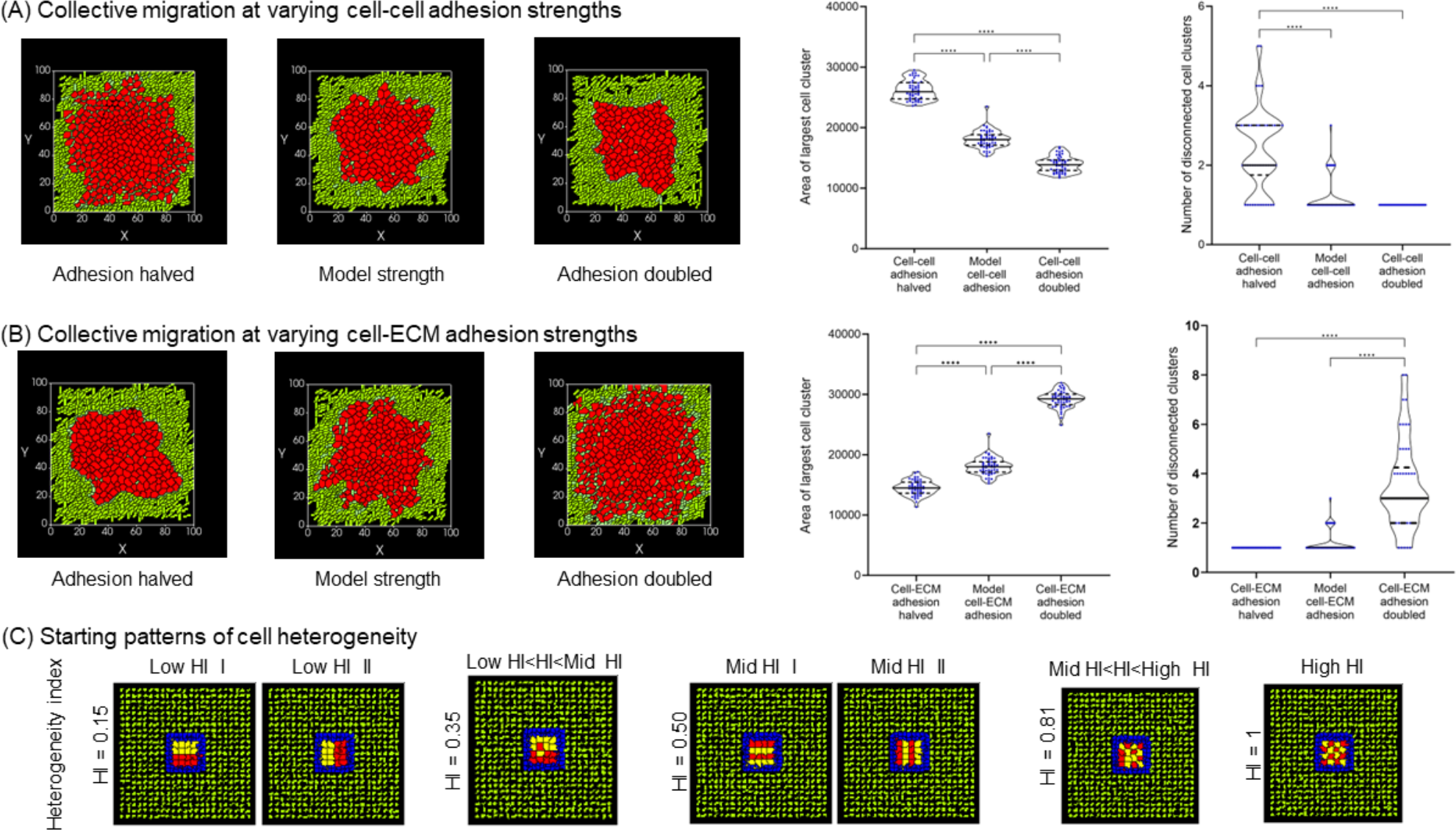
Introduction to the computational framework for the study of regulation of collective cancer migration by spatial cell patterns with varying cellular and matrix adhesion. (A) Phenotypes of tumor populations of cells (in red) at the end of simulations run (1200 MCS), wherein cells had model cell-cell adhesion strength (that had been shown to give rise to collective migration in Pramanik et al., J. Theor. Biol., 2021; center), with half the model cell-cell adhesion strength (left) and with double the model cell-cell adhesion strength (right). Violin plots with medians for area of largest cell cluster (used as a measure of collective migration) and number of disconnected cell clusters (used as a measure of dispersed migration) for 50 simulations each for the three cell-cell adhesion strengths are shown in right. (B) Phenotypes of tumor populations of cells (in red) at the end of simulations run (1200 MCS), wherein cells had model cell-ECM adhesion strength (that had been shown to give rise to collective migration in Pramanik et al., J. Theor. Biol., 2021; center), with half the cell-ECM adhesion strength (left) and with double the cell-ECM adhesion strength (right). Violin plot with medians for area of largest cell cluster (used as a measure of collective migration) and number of disconnected cell clusters (used as a measure of dispersed migration) for 50 simulations each for the three cell-ECM adhesion strengths are shown in right. (C) Phenotypes of tumor populations at the beginning of simulations comprising two types of cancer cells (colored yellow and red) with distinct cell-cell or cell-ECM adhesion strengths, arranged in spatial patterns with an increasing degree of heterogeneity index (see appropriate Methods section for definition) from left to right (leftmost: consecutive horizontal and vertical patterns with HI = 0.15; consecutive pattern with a single cell pair swap with HI = 0.35; alternate horizontal and vertical patterns with HI = 0.50; pattern with a single cell pair swap with HI = 0.81; checkered pattern with HI = 1) Statistical significance for the measurements in the simulations computed using one-way ANOVA with Tukey Kramer post hoc comparisons.

We subsequently moved to heterogeneous pattern cases, wherein the above-mentioned adhesion strengths can be observed to spatially vary across a cell population (in this paper, red- and yellow-colored cells represent cells with distinct adhesion strengths). We quantified this variation through the heterogeneity index (HI), which measures the mean of immediate dissimilar cell neighbors for each cell in the starting cell population **(Fig 1C)**. In order to explore the effects on migration across the heterogeneity index, we examined distinct spatial patterns, which we call low HI I and II (here, the adhesion strengths have been kept the same for two consecutive rows of red or yellow cells resembling a tumor with two distinct well-sorted cell populations)); mid HI I and II (here, the adhesion strengths alternate between rows of red and yellow cells resembling a tumor with two distinct but moderately intermingled cell populations); and finally a high HI pattern (here, the adhesion strength alternates between two consecutive cells within the population resembling a tumor with prolific cell state mingling). The rationale for examining the consecutive or alternating arrangements along two axes, and hence two versions each for low and med HI) was to discount any confounding effects by the alignment of fibrillar ECM (from bottom-left to top-right of the simulation frame) on the symmetry of the migrating cell pattern. In addition to these symmetric patterns, we also swapped a pair of red and yellow cells each for the low HI and high HI case to evince asymmetric patterns whose HI therefore lies intermediate between low and mid HI patterns, and mid HI and high HI patterns, respectively.

To examine the role of heterogeneity in cell-cell adhesion in CCM, we first carried out simulations on progressively intermingled populations, wherein the two cellular subsets showed adhesion to their own respective subset cell type that were half and double the model adhesion strength (i.e., yellow cells having cell-cell adhesion half of model cell-cell adhesion and red cells having cell-cell adhesion double of model cell-cell adhesion). In addition, for each simulation we varied the inter-subset cell adhesion strength from half to double the model cell-cell adhesion strength. We observed that in the context of lower inter-subset adhesion strength **(Fig 2A)**, i.e., when the two populations were least adhered to each other, the CCM shown by patterns increased in direct proportion with their HI **(Fig 2B**; ANOVA p<0.0001, significant differences observed between patterns with widely diverged HI through post hoc comparisons). Observing the population phenotypes at the end of simulations showed that low inter-subset adhesion strength ensured that irrespective of how intermingled they were at the beginning of the simulation, the subsets eventually sorted out from each other. None of the patterns under such contexts showed any notable dispersion (**Fig 2C**).

**Figure 2:**
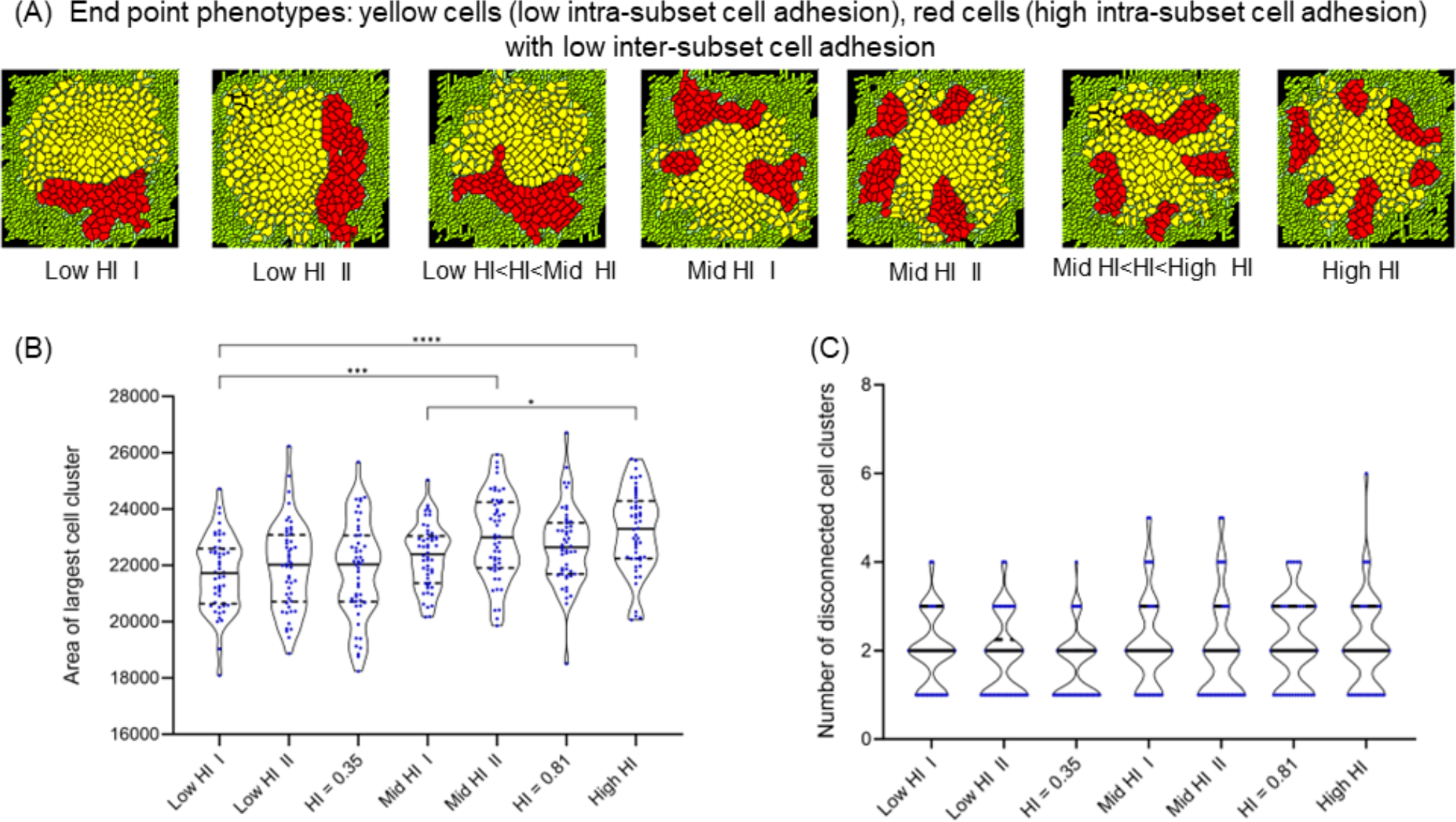
Collective migration in tumor populations with heterogeneous intra-subset cell adhesion and low inter-subset cell adhesion. (A) Endpoint (1200 MCS) phenotypes of tumor populations with progressively increasing HI values of starting patterns with yellow cells (low intra-subset adhesion), red cells (high intra-subset adhesion) and low inter-subset adhesion. (B) Violin plot with median of area of largest cell cluster (used as a measure of collective cell migration) for 50 simulations each for patterns from low-to high HI values. (C) Violin plot with median of the number of disconnected cell clusters (used as a measure of dispersed migration) for 50 simulations each for patterns from low-to high-HI values. Statistical significance for the measurements in the simulations computed using one-way ANOVA with Tukey Kramer post hoc comparisons.

The relationship between CCM and HI switched when inter-subset adhesion strength was assigned a high value **(Fig 3A)**, i.e., when the two populations strongly adhered to each other: here, migration of patterns showed an inversely proportional relationship with their HI **(Fig 3B;** ANOVA p<0.0001, significant differences observed between patterns with widely diverged HI through post hoc comparisons). In addition, we observed that the high inter-subset adhesion strength engendered strong intermingling of subsets at the end of simulations carried out for populations with varying degree of HI. This implies that adhesion-heterogeneous tumor populations that are well intermingled will collectively migrate better only in the context of weak inter-niche adhesion, i.e., when heterotypic cell-cell adhesion is sparse. Under strong inter-niche adhesive contexts, coarsely intermingled tumor populations migrate better. As in Fig 2C, there was very little DCM shown by such migrating cell populations **(Fig 3C)**.

**Figure 3:**
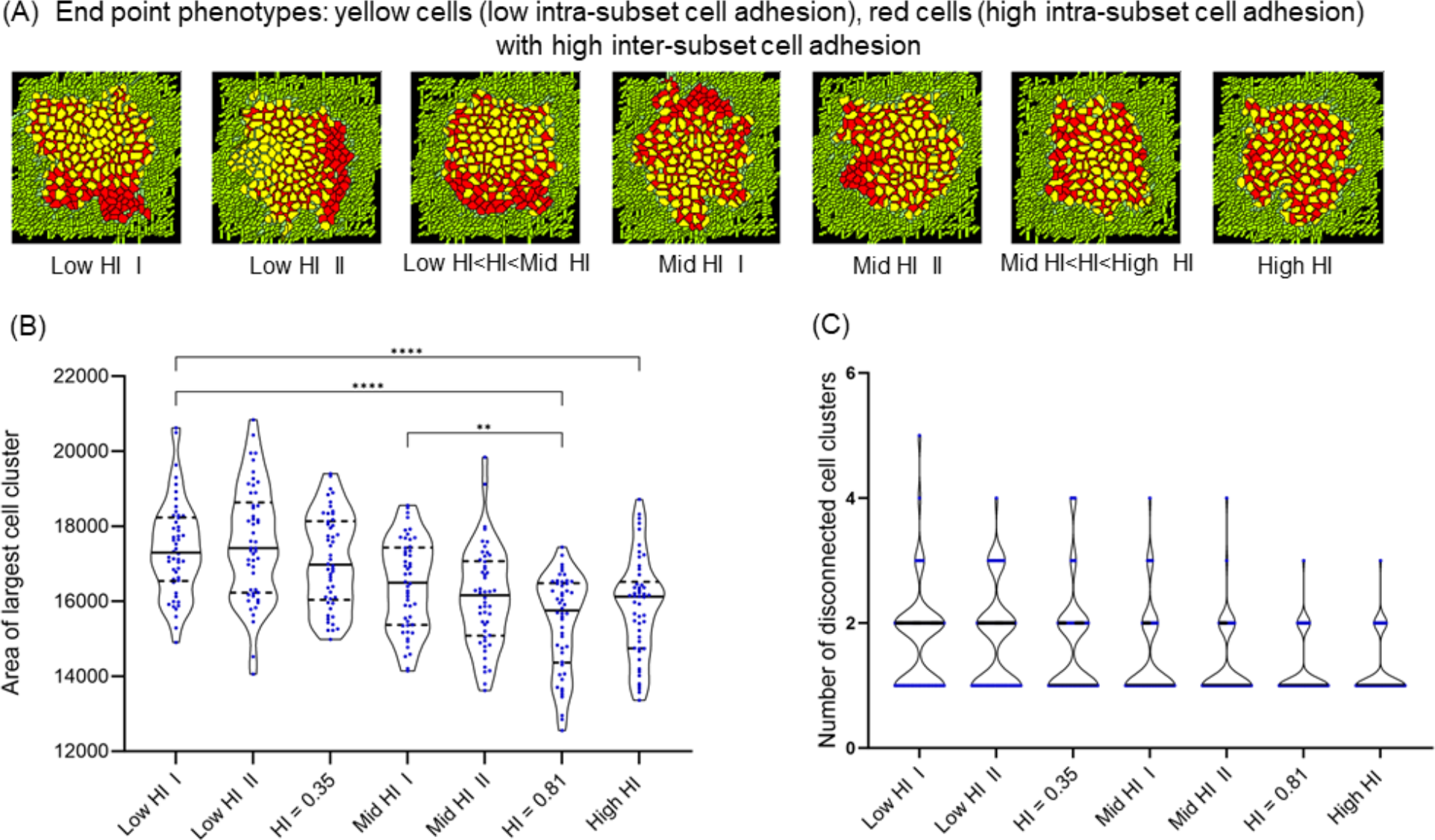
Collective migration in tumor populations with heterogeneous intra-subset cell adhesion and high inter-subset cell adhesion. (A) Endpoint (1200 MCS) phenotypes of tumor populations with progressively increasing HI values of starting patterns with yellow cells (low intra-subset adhesion), red cells (high intra-subset adhesion) and high inter-subset adhesion. (B) Violin plot with median of the area of largest cell cluster (used as a measure of collective cell migration) for 50 simulations each for patterns from low-, to high-HI values. (C) Violin plot with median of the number of disconnected cell clusters (used as a measure of dispersed migration) for 50 simulations each for patterns from low-to high-HI values. Statistical significance for the measurements in the simulations computed using one-way ANOVA with Tukey Kramer post hoc comparisons.

### Influence of heterogeneous cell-ECM adhesion on the collective migration phenotype

We next investigated the migration of heterogeneous cell patterns under different cell-ECM adhesion strengths **(Fig 4A)**, while considering no heterogeneity in cell-cell adhesion. For simulations in which the two subpopulations showed halved and doubled model cell-ECM adhesion strengths respectively (i.e., yellow cells having cell-ECM adhesion half of model cell-ECM adhesion and red cells having cell-ECM adhesion double of model cell-ECM adhesion), we found a progressive increase in CCM as the HI in cell-ECM adhesion strength was increased **(Fig 4B**; ANOVA p<0.0001, significant differences observed between patterns with widely diverged HI through post hoc comparisons). We observed only insignificant variations in DCM **(Fig 4C)**. The population phenotypes at the end of the simulations showed that the intermingling of subsets was broadly recapitulative of their starting pattern **(Fig 4A)**.

**Figure 4:**
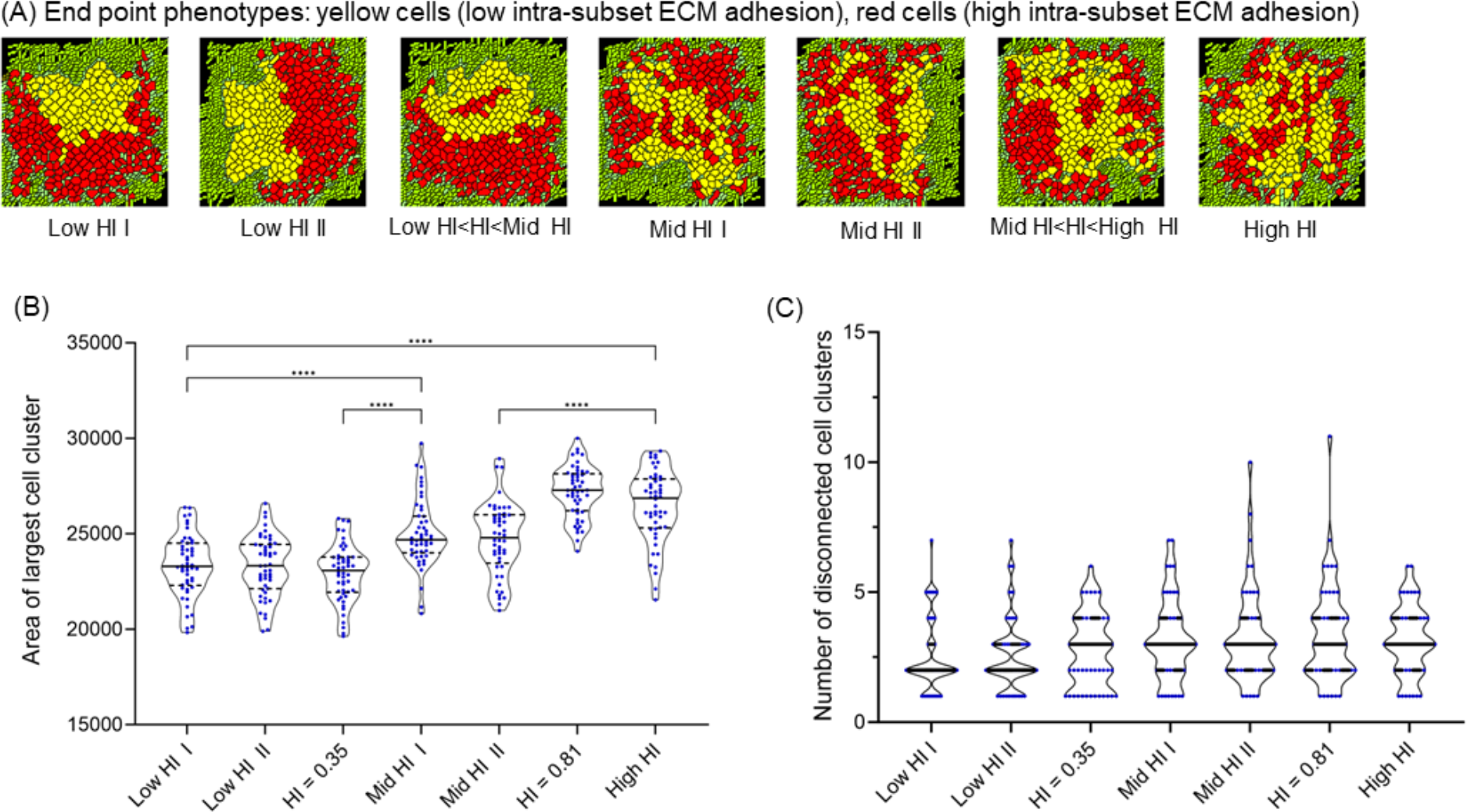
Collective migration in tumor populations with heterogeneous intra-subset cell-ECM adhesion. (A) Endpoint (1200 MCS) phenotypes of tumor populations with progressively increasing HI values of starting patterns with yellow cells (low intra-subset cell-ECM adhesion), red cells (high intra-subset cell-ECM adhesion). (B) Violin plot with median of the area of largest cell cluster (used as a measure of collective cell migration) for 50 simulations each for patterns from low-to high-HI values. (C) Violin plot with median of the number of disconnected cell clusters (used as a measure of dispersed migration) for 50 simulations each for patterns from low-to high-HI values. Statistical significance for the measurements in the simulations computed using one-way ANOVA with Tukey Kramer post hoc comparisons.

### Incorporating heterogeneity in both cell-cell and cell-ECM adhesion components

So far, we examined the effects of heterogeneity in adhesion strengths of cells with each other or with their surrounding ECM separately from each other. However, signaling alterations associated with tumorigenesis cross-link such traits allowing them to combinedly shape phenotypes. For example, epithelial to mesenchymal transition (EMT) seen in cancer cells is associated both with a depletion in cell-cell adhesion and an increase in matrix adhesion [24], [25]. How do such associative changes in adhesion strengths regulate migration across a tumor population? To answer this question, we first carried out simulations, wherein CCM and DCM were assessed in homogeneous cellular populations with variable levels of both adhesion types **(Fig 5A)**. Unsurprisingly, cells with stronger cell-cell adhesion but weaker cell-ECM adhesion (double and half of the model adhesion strengths, respectively; resembling stereotypically strong epithelial states) exhibited poor CCM **(Fig 5B**; ANOVA p<0.0001, significant differences observed between patterns through post hoc comparisons). On the other hand, populations with weaker cell-cell adhesion and stronger ECM adhesion (resembling stereotypically strong mesenchymal states), collectively migrated further but also dispersed into the surrounding ECM as discrete clusters **(Fig 5B-C**; In Fig 5C, ANOVA p<0.0001, significant differences observed between mesenchymal pattern with control and epithelial patterns assessed through post hoc comparisons) as has been observed in experiments [26].

**Figure 5:**
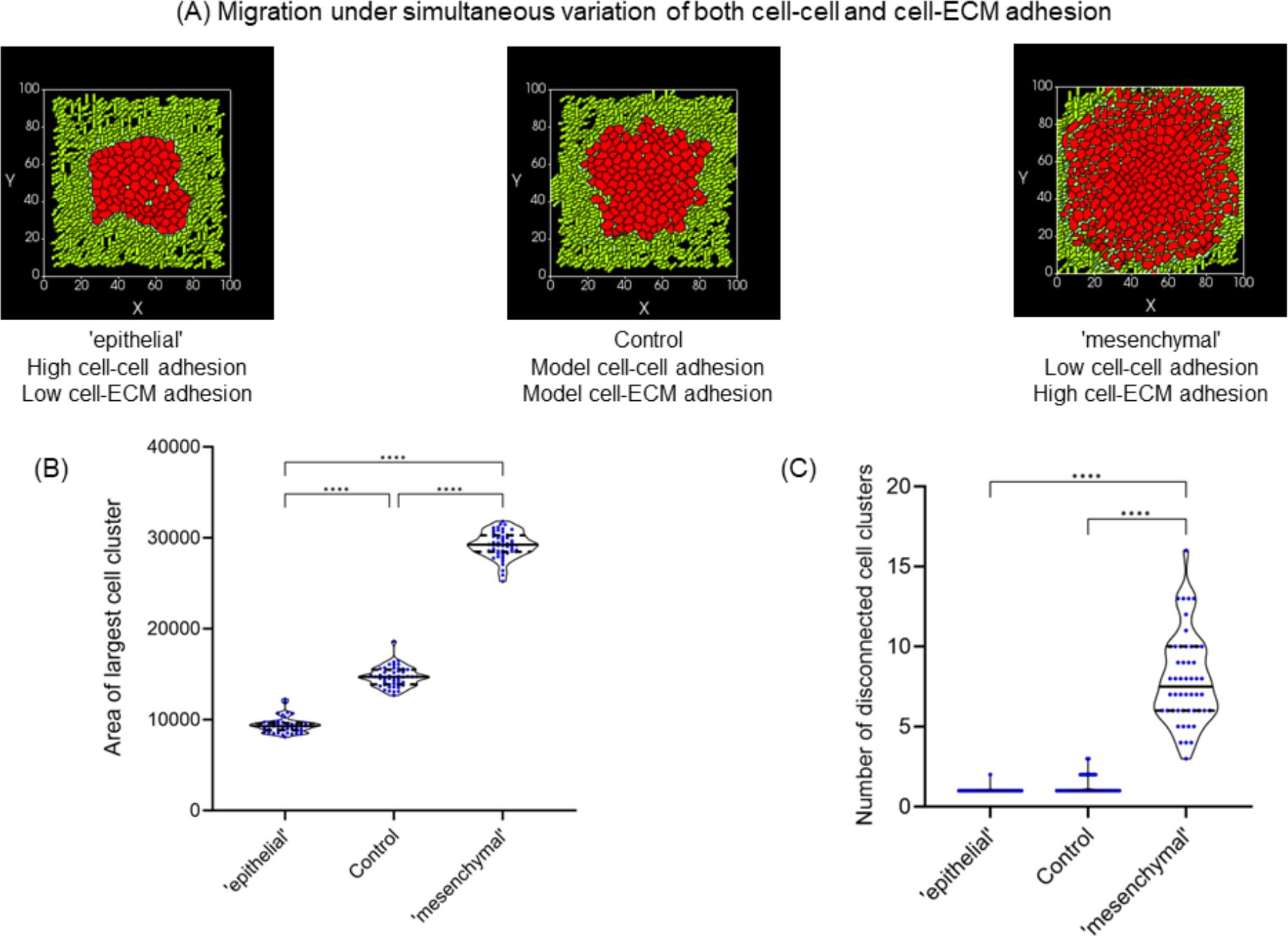
Migration upon simultaneous variation of both cell-cell and cell-ECM adhesion strengths. (A) Phenotypes of tumor populations of cells (in red) at the end of simulations run (1000 MCS), wherein cells had model cell-cell and -ECM adhesion strength (center; control), with double of model cell-cell and half of model cell-ECM adhesion strength (left; ‘epithelial’), and with half of model cell-cell and double of model cell-ECM adhesion strength (right; ‘mesenchymal’). (B) Violin plot with median of area of largest cell cluster for ‘epithelial’, control (collective migration) and ‘mesenchymal’ (multimodal mode of migration). (C) Violin plot with median of number of disconnected cell clusters for ‘epithelial’, control and ‘mesenchymal’. Statistical significance for the measurements in the simulations computed using one-way ANOVA with Tukey Kramer post hoc comparisons.

Heterogeneity in cellular and histological phenotypes driven by non-genetic, genetic, and microenvironmental cues is the norm of tumor populations [7], [27]. Among the axes of heterogeneity, there is mounting evidence for the concurrent presence within cancer populations of cellular phenotypes across the epithelial to mesenchymal spectrum [28], [29]. To test the consequences of the spatial arrangement of a heterogeneous population of epithelial and mesenchymal cells on their migration, we conducted simulations of patterns with different HI values and with high inter-subset cell adhesion strength **(Fig 6A)**. We observed an inversely proportional relationship between HI values and the extent of CCM **(Fig 6A** left graph; ANOVA p<0.0001, significant differences observed between patterns with widely diverged HI through post hoc comparisons). Moreover, regardless of HI values, all end point simulations showed a strong intermingling of epithelial and mesenchymal cells. In contrast, when inter-subset cell adhesion strength is kept low, HI values show a directly proportional relationship with the extent of CCM **(Fig 6B** left graph; ANOVA p<0.0001, significant differences observed between patterns with diverged HI through post hoc comparisons). In these simulations, regardless of HI values, the two populations got sorted out efficiently. For high inter-subset adhesion strengths, post hoc comparisons showed little difference in DCM between different HI patterns. However, a progressive increase in DCM was observed in the low inter-subset cell adhesion case, where high HI patterns showed the strongest dispersion compared with all other patterns (ANOVA p<0.0001, significant differences observed between patterns with diverged HI through post hoc comparisons).

**Figure 6:**
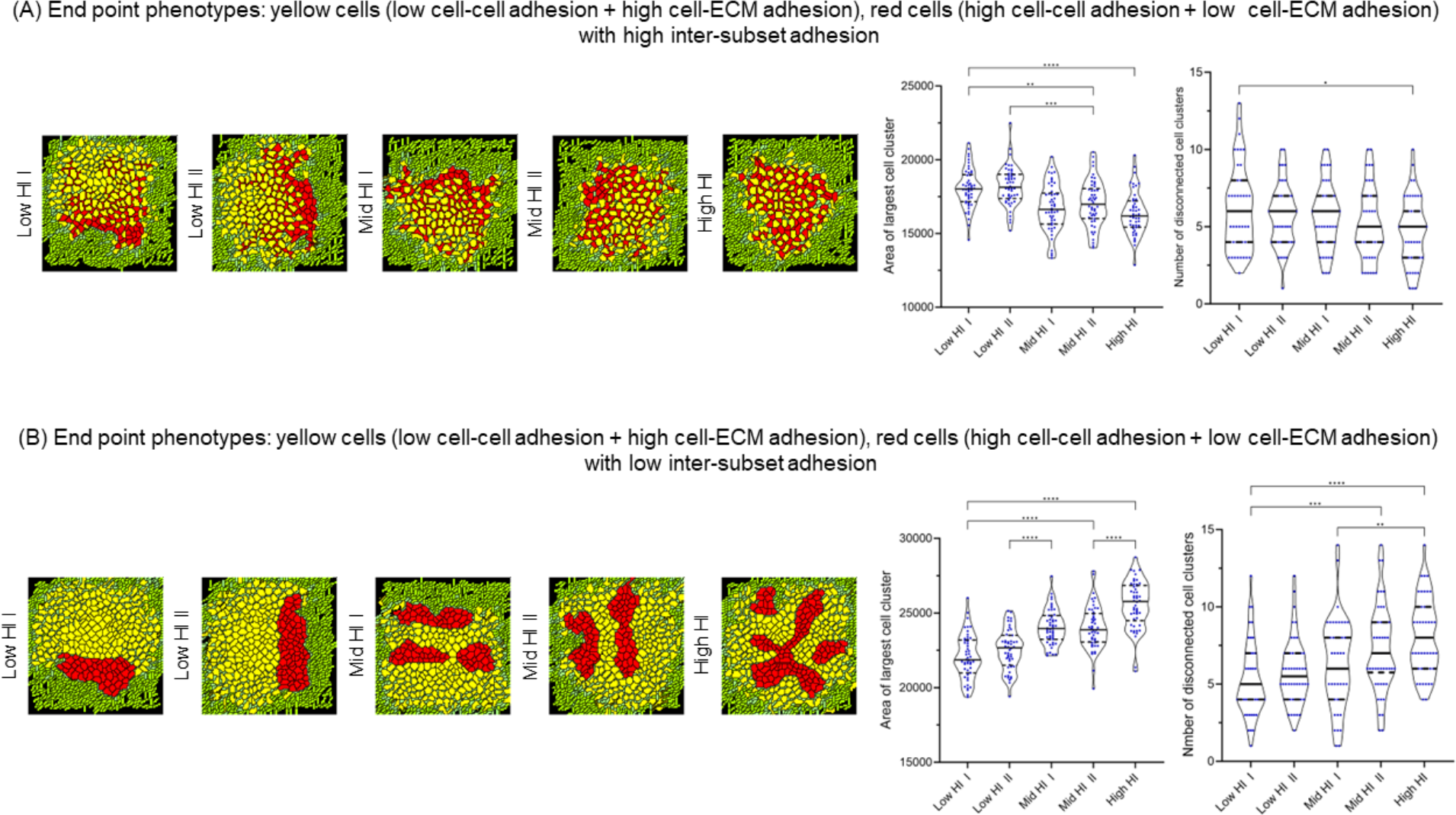
Migration in tumor populations with heterogeneity in intra-subset cell adhesion and cell-ECM adhesion. (A) Endpoint phenotypes of tumor populations with progressively increasing HI values of starting patterns with yellow cells (low intra-subset cell adhesion and high cell-ECM adhesion), red cells (high intra-subset cell adhesion and low cell-ECM adhesion) with high inter-subset cell adhesion. Violin plots with medians for area of largest cell cluster and number of disconnected cell clusters for low-, mid- and high hi value spatial patterns on right. (B) Endpoint phenotypes of digital tumor populations with progressively increasing HI values of starting patterns with yellow cells (low intra-subset cell adhesion and high cell-ECM adhesion), red cells (high intra-subset cell adhesion and low cell-ECM adhesion) with low inter-subset cell adhesion. Violin plots with medians for area of largest cell cluster and number of disconnected cell clusters for low-, mid- and high-hi value spatial patterns on right. Statistical significance for the measurements in the simulations computed using one-way ANOVA with Tukey Kramer post hoc comparisons.

We next asked if assessing the HI of initial tumor population pattern is sufficient to predict the relative extent of collective migration in heterogeneous cancer cell populations. To our surprise, we observed patterns with relatively lower HI (for cell-ECM heterogeneous adhesion case), which showed higher CCM **(Fig 7;** Student’s t-test, p=0.005) but insignificantly different DCM **(Fig 7**; Student’s t-test, p=0.08). These patterns were characterized by a relatively greater partitioning of cells with higher adhesion to collagen to the radially outer edge of the tumor population. Such patterns have especially been shown to be associated with high migrative efficiency in experiments with cocultures of aggressive breast cancer cells comprising niches that are distinct in their cell-ECM strengths [30]. Our observations therefore suggest that cancer cells with greater ability to adhere to stromal fibrillar collagens contribute further to CCM when they are spatially well-distributed across the tumor, but especially to the outer edge, where they can exert their adhesive effects on the neighboring ECM more efficiently than other patterns.

**Figure 7:**
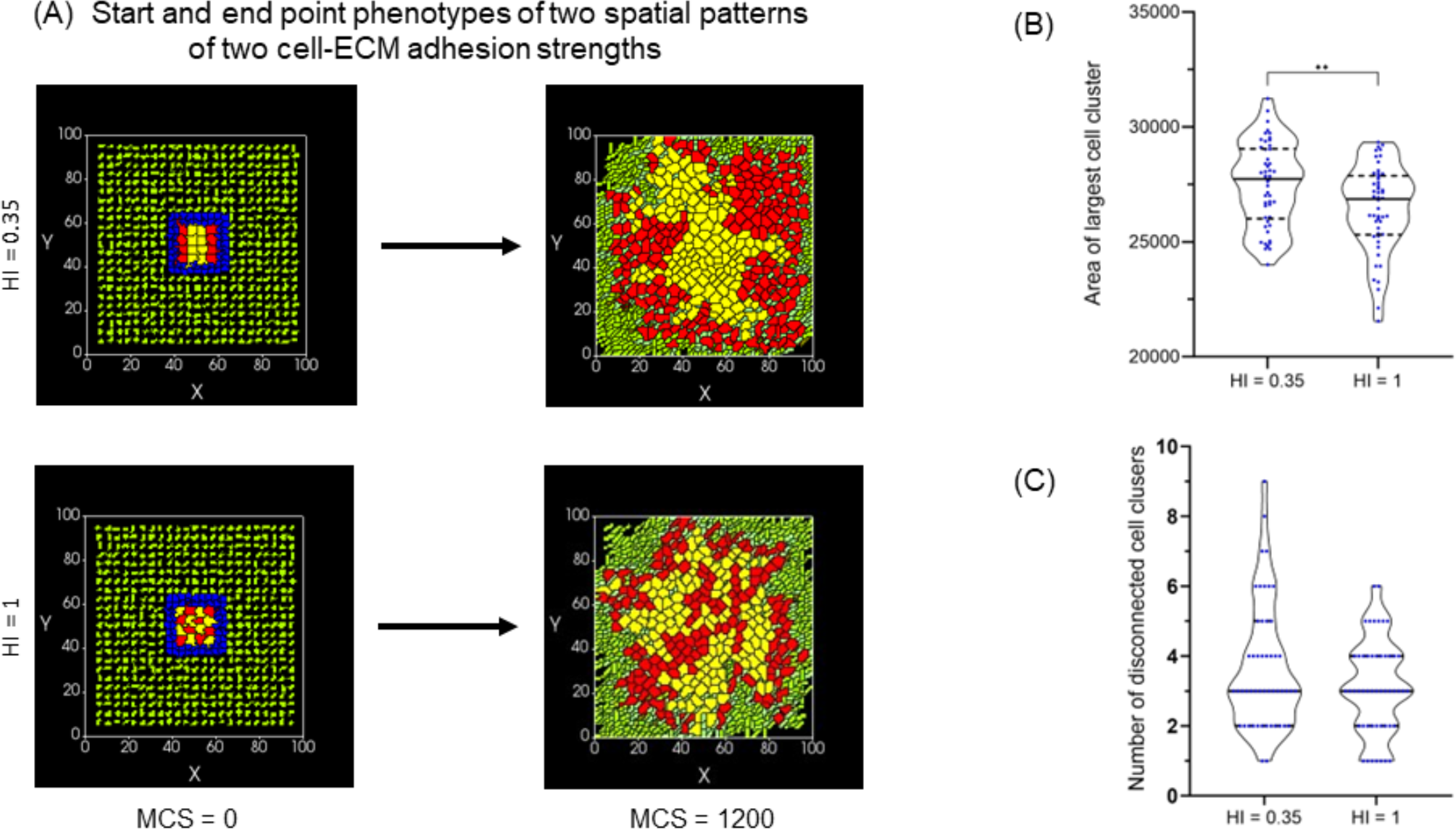
A special case of a starting spatial pattern with HI smaller than High HI showing higher migration for cell-ECM adhesion strength variation. (A) Start and end point phenotypes of two spatial patterns with cell-ECM adhesion being double of model cell-ECM adhesion for red cells and half for yellow cells. (B) Violin plot with median of area of largest cell cluster for spatial patterns with HI = 0.35 and HI = 1. (C) Violin plot with median of number of disconnected cell clusters for spatial patterns with HI = 0.35 and HI = 1. Statistical significance for measurements in the simulations computed using two-tailed unpaired Student’s t-test.

## Discussion

Tumors show heterogeneous signatures across molecular and cellular scales. Clonal populations may exist with distinct single nucleotide and copy number variations (SNVs and CNVs, respectively) [31], protein expression [32], cellular phenotypes [33], and histopathological features [34]. However, the extent to which distinct heterogeneous behaviors correspond spatially across scales is yet to be well understood (although see [34] for a demonstration of a correspondence between variation in nuclear morphology and genomic instability). Various reports on spatial heterogeneity in primary tumors report a gradient of EMT such that the leading edge is relatively more mesenchymal [35]–[37]. Such observations have been explained by computational models invoking a diffusible EMT-inducing signal such as TGFβ, known to be secreted by stromal cells [38], however, these studies do not investigate the functional contributions of spatial heterogeneity in tumor progression. Further, *in vitro* studies on collective cell migration highlight that this EMT gradient is rather dynamic, given the exchange of leader and follower cells in a coordinated manner [39], [40]. Thus, elucidating the functional roles of spatial heterogeneity becomes crucial.

While a series of elegant studies show spatial heterogeneity in clonal populations as confounding management and potentiating metastasis and mortality [41]–[43], they do not explicitly study the consequences of the degree of intermingling between two or more heterogeneous signature-expressing clones. The reason for this could be the difficulty in inferring cell behavioral traits from (multi)omic expressions assayed through sequencing studies. Studying lung adenocarcinoma, Wu and workers have broken fresh ground by demonstrating two distinct diversification patterns, which they call clustered and random geographic diversification patterns [44]. These two correspond broadly to our two ends in the spectrum of heterogeneity index patterns. They show that the random pattern was more deleterious in its association with disease recurrence and mortality. Our simulations provide a formal and contextual basis for such observation. It also allows us to propose that the association of such a random diversification pattern with cancer cell migration is facilitated under conditions of lower cell-cell adhesion between heterogeneous niches. Under higher adhesion conditions, a random pattern may have opposite consequences.

Recent studies on primary tumor explants show heterogeneity in rigidity across multicellular and cellular scales, with soft motile cells surrounding islands of immotile stiff tumor epithelia. The relation between cell-cell adhesion and rheological behaviors of tissues and tumors is driven by reciprocal effects of substrata rigidity and organization and expression of cell adhesion proteins and cytoskeletal regulators [45], [46]. Under non-compliant confining conditions, cancer cells have been observed to show an elevation in cadherin presentation on the cell surface [47], which induces proliferation in such populations. Whether this may influence the intermingling between soft and rigid tumor cells remains unclear. Although our simulations do not impose rheological properties onto cells, we do observe that the patterning, or the intercellular arrangement of cells sufficiently unjams the cells under specific conditions. Especially under conditions of low inter-niche adhesion, or under conditions wherein niches with greater ECM adhesion are broadly distributed across the exterior of the tumor, we see higher unjamming behavior. What was striking in our simulations was that changing either cell-ECM or cell-cell adhesion separately increased migration but quantitatively, while retaining the collective nature. However, combinatorial alterations led to a qualitative shift in behavior from collective to dispersed (within the surrounding ECM) behavior. This potentially explains why the transition from epithelial character to mesenchymal requires cells to cross-regulate cytoskeletal elements that mediate adhesion to ECM substrata while also depleting cell-cell adhesion mediators such as E-cadherin.

Despite the predictive power of our framework, we would like to highlight the latter’s limitations. Our assays are still in a two-dimensional computational space, due mostly to the restraint in computational power required to have a three-dimensional ECM fiber network. Our framework is also not amenable to the direct incorporation of mechanical properties that play an important role in cell-ECM interactions. Third, the consequences of genomic and transcriptional heterogeneity are manifested not only in terms of cellular adhesion but also in traits such as migration, ECM degradation, and proliferation, which may be modulated in association with or independently of the former. Future iterations of the model will incorporate signaling networks that link multiple traits and which may be amenable to variation in signaling strengths indicative of the consequences of genetic and epigenetic aberrations. Our investigations in this manuscript examined heterogeneity in adhesion within the parametric constraints of collective cell migration. Our next effort will be focused on examining the effects of spatial variation in adhesion under initial conditions of dispersed or multimodal migrations.

We conclude by highlighting the broader implications of our study: although our model is motivated in its assumptions and histopathological foundations by tumorigenic context, the framework itself is as easily adaptable to embryological problems wherein collective cell migration has been found to play a role in several organogenetic systems such as glandular morphogenesis, gastrulation, somitogenesis, and limb development. There is burgeoning evidence through single-cell sequencing for the presence of cellular heterogeneity within developing tissues and organs during embryogenesis: in fact, polyclonal modes of embryogenesis have been reported across diverse phyla [48]. Our results suggest that adhesion heterogeneity allows a cellular population to migrate effectively while also retaining evolvability towards switching between multicellular to unicellular phenotypes. A translational ramification of our study is that interactions based on differential adhesions can mediate qualitative shifts in the ‘graininess’ of niche heterogeneity in growing tumoroid populations. Cellular masses that start out with finely intermingled niches may show niche-sorting during their progression: biopsying such tumors at a single spatial locus may lead to an underestimation of intratumoral heterogeneity. Our simulations suggest a multi-temporal multi-locus strategy for the assessment of heterogeneity within malignant tumors. Although not addressed in this manuscript, it is pertinent to ask what may lead to the manifestation of the heterogeneous patterns we have worked with in this paper. Whereas low HI patterns are phenomenologically indicative of differentially sorted cells, evidenced in tissues with altered adhesion molecules, high HI index patterns are suggestive of operational mesoscale signaling of juxtracrine or paracrine nature, such as Notch-Delta signaling [49]. The interplay between the latter and adhesion will be probed in future studies.

## Data availability

The code for the manuscript can be accessed at https://github.com/cvsp-res/spatial_hetero.git

## Authors’ contribution

RB and MKJ framed the problem statement. CVSP carried out the simulations. RB, MKJ, and CVSP analyzed the results of the simulations. RB, MKJ, and CVSP wrote and edited the manuscript.

## Materials & Methods

### Modelling framework and CompuCell3D

CompuCell3D is an agent-based simulation framework that is developed to computationally model systems of cell-, developmental- and cancer biology. It is primarily based on the Cellular Potts model (CPM) / Glazier-Graner-Hogeweg (GGH) model [50]. CompuCell3D (CC3D) combines CPM/GGH with partial differential equation (PDE) solvers for chemical fields and other models for spatiotemporal modeling by defining spatially extended generalized cells, which can represent single biological cells, their clusters or even sub-compartments and subdomains of non-cellular materials [51]

The CPM hamiltonian/effective energy, the cornerstone of all CC3D simulations describes cell behaviors and interactions by incorporating contact energy terms, which determine the extent of interaction between different cell types in the simulations. Positive contact energy values indicate the extent of repulsion between different entities in the simulation. In the context of biology, it is the adhesion between the entities which has significance and implications. The relation between adhesion and contact energy in CC3D is inverse, i.e., adhesion is negative of contact energy.

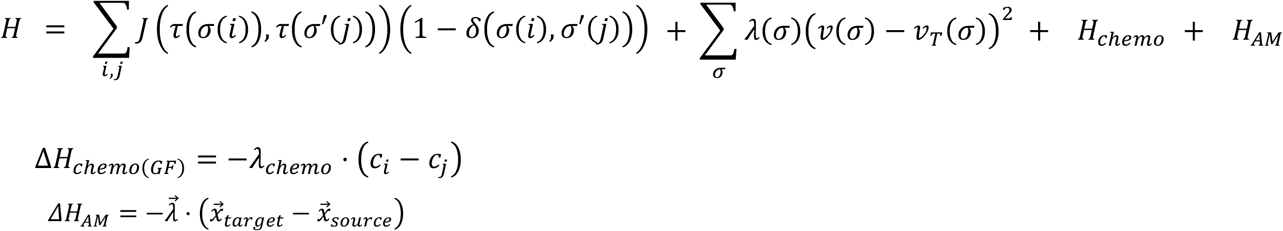

The time evolution of simulations in CC3D involves many attempts to copy cell indices between neighboring pixels. On a successful index copy attempt, the volume of the source cell goes up while that of the target cell goes down by one pixel (Please see the Introduction to Compucell3D manual).

A Monte Carlo Step (MCS) consists of one index copy attempt for each pixel in the cell lattice. Modified Metropolis with Boltzmann acceptance function regulates attempts to copy indices. At any given MCS, if the index copy attempt is successful, the effective energy of the new configuration will be evaluated. If the change in effective energy is negative, then that index copy attempt is considered and implemented which will be depicted in the next MCS in the simulation. If the change in effective energy is positive, then there is a probability associated with that index copy attempt to happen described by the Boltzmann acceptance function.

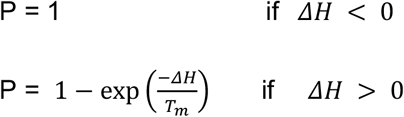

where ΔH is change in effective energy if the copy occurs and *T*_0_ is a parameter describing the amplitude of cell-membrane fluctuations [51]. The first term of effective energy corresponds to contact energies and adhesion. The second term in effective energy is the volume constraint energy term, which is analogous to the spring energy term. It ensures the cell volume 𝒱 (σ) does not deviate from the cell’s target volume 𝒱 _*T*_(σ). λ (σ) is like a spring constant that denotes the inverse compressibility of the cell. 𝒱 (σ) of each cancer cell is governed by growth equation [3]. Δ*H*_*chemo(GF)*_ and *H*_*AM*_ terms of the current model are specific to cancer cells only in the simulations. Δ*H*_*chemo(GF)*_ enables chemotaxis of cancer cells that represents a biased motility of cells in response to the growth factor chemical gradient they are exposed to. *c*_6_ represents the concentration of the chemical field (GF) at the index-copy target pixel (*i*) while *c*_7_ corresponds to the same at the index-copy source pixel (*j*). *λ* _*chemo*_ is the strength of the chemotaxis that cells respond to. Δ*H*_*AM*_ is used to model the forces between the cells. (AM in the subscript corresponds to active motility of the cells). 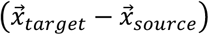 is the difference between position vectors of source and target pixels while 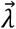 is the strength and direction of external force between the cells. In the current model, Δ*H*_*AM*_ it is used to take into consideration the forces between the cells that give rise to random cell motion. The respective steppable (CellMotility steppable) provides an external force on the center of mass of the cells which changes direction randomly every MCS.

### Simulation details of the model

The domain of simulation we constructed is a 100 x 100 x 1 square lattice with a non-periodic boundary condition consisting of seven different entities that are explained below [3]. In the simulations, the lengths and volume dimensions are measured in terms of number of pixels or voxels and the Monte Carlo step (MCS) is the unit of time.

#### Medium

These are unassigned cell types in the background on which the computational model is constructed in the simulation domain.

#### Cancer cells

These cells are positioned at the center of the simulation surrounded by basement membrane/laminin. There are two types of cancer cells: Cancer cell subset - 1 (C1) which is indicated by red color and Cancer cell subset -2 (C2) which is indicated by yellow color. Each cancer cell has a volume of (4×4×1) pixels, and the tumor is a grid of 16 such cancer cells with 8 cells of each type. Only the cancer cells in the simulation have the property to respond to chemical gradients of growth factor, grow, proliferate, and degrade ECM.

#### BM (Laminin-rich ECM)

This cell type corresponds to the thick basal lamina which surrounds and holds together the luminal epithelial cells in the mammary duct. In simulations, this is represented by a two-layered, blue-colored annular basement membrane binding the cancer cells together and separating them from collagen1/ECM [3].

#### Collagen I (Co1)

These cells mimic the fibrillar extracellular matrix (ECM) surrounding the tumor. Cancer cells degrade Collagen I and migrate through it.

#### C_lysed

This cell type is used in an intermediate step during matrix degradation and regeneration. The dynamics of reaction-diffusion of the chemicals secreted by cancer cells allow for the degradation of BM and Co1 matrix cells. Upon meeting certain criteria for degradation, the BM or Co1 cell type of a particular cell becomes C_lysed, although retaining the shape and size of the cell. These cells track the MCS from their individual degradation event and transform it into newly synthesized matrix cells after 20 MCS.

#### NC1

This cell type is intended to mimic the ‘cancer matrisome’, the newly synthesized matrix cells are denoted as NC1. These cells are almost identical to Collagen I in their behavior and can undergo further degradation to become C_lysed and subsequently after 20 MCS, would become NC1 again.

#### Reaction-Diffusion, growth, and proliferation equations

Although our multiscale model deploys reaction-diffusion-based interactive dynamics of soluble matrix metalloproteinases and their inhibitors, this has not been analyzed formally in this manuscript and readers are encouraged to refer to our previous paper for further details on this aspect of the model description [3].

### Starting spatial patterns and heterogeneity

The spatial heterogeneity in tumors at the beginning of the simulation was represented through the following five spatial patterns: namely Low HI I, Low HI II, Mid HI I, Mid HI II and High HI. Besides them, spatial patterns intermediate to Low- and Mid-HI, intermediate to Mid- and High-HI, and a special case (**Fig 7**) were considered.

Spatial heterogeneity at the cellular level is considered as the ratio of number of dissimilar cells surrounding the given cell to the total number of cells surrounding it. Summing up these ratios calculated for all cells of one subtype in each spatial pattern yields the heterogeneity index (HI) of that cell subtype. HI of both cell subtypes is considered as HI of the spatial pattern. In the case of unequal HI between cell subtypes, the average of both the values is taken as HI of the spatial pattern. In the current definition of HI only 1^st^ order neighbors i.e., adjacent cells that share a common edge are considered as surrounding cells. Therefore, it shall be 4 for an interior cell and 3 for a peripheral cell except the corner cells. The corner cells will have 2 surrounding cells.

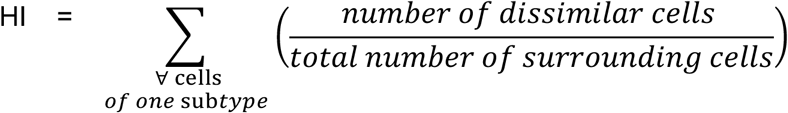

The HI calculations for red cells in spatial patterns in (**Fig 1C**) starting from topmost left and traversing clockwise are shown below.

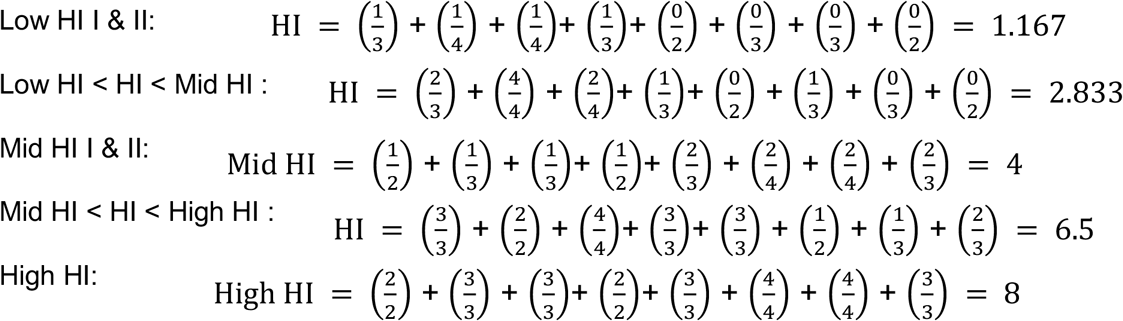

Scaling the above values of HI from 0 to 1, we get the new HI values as

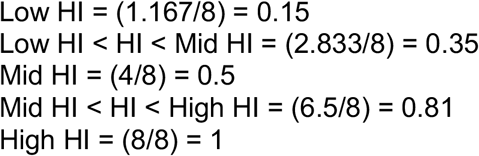

Scaled HI value for specific case pattern = (0.4167 + 0.29167)/2 = 0.35 where 0.4167 is HI of red subset of cells and 0.29167 is HI of yellow subset of cells.

### Statistical Analysis

All Compucell3D simulation screenshots at 1200 MCS time step were taken (1000 MCS for simultaneous variation of both cell-cell and cell-ECM case) to do image quantification analysis in MATLAB. Considering that the cellular dynamics incorporated in our model are stochastic in nature and to ensure a large sample size for statistical analysis, 50 replicates of the cases were carried out. For statistical analysis, one-way ANOVA was performed followed by the Tukey-Kramer test as a post-hoc analysis in GraphPad on the heterogeneous and homogeneous simulations of every case to draw inferences and conclusions. Significance (p value) is represented as *, where * ≤ 0.05, ** ≤ 0.01, *** ≤ 0.001, and **** < 0.0001.

